# Quantitative classification of energy landscapes inferred from single nanoparticle tracking of membrane receptors inside nanodomains reveals confinement functional and molecular features

**DOI:** 10.1101/2023.02.13.528279

**Authors:** Chao Yu, Maximilian Richly, Thi Thuy Hoang, Mohammed El Beheiry, Silvan Türkcan, Jean-Baptiste Masson, Antigoni Alexandrou, Cedric I. Bouzigues

**Affiliations:** Laboratoire Optique et Biosciences, CNRS UMR74645, Inserm U1182, Ecole Polytechnique, Institut Polytechnique Paris, Palaiseau, France; Decision and Bayesian Computation, Computational & Neuroscience departments, CNRS UMR 3751, Institut Pasteur, Université Paris Cité, Paris, France

## Abstract

The cell membrane organization has been hypothesized for a long time to have an essential functional role, through the control of membrane receptor confinement in micro- or nanodomains. Several mechanisms have been proposed to account for these properties, though some features of the resulting organization have remained controversial, notably the nature, size, and stability of cholesterol- and sphingolipid-rich domains called rafts.

Here, we quantitatively probed the energy landscape experienced by single nanoparticle-labeled membrane receptors - epidermal growth factor receptors (EGFR), transferrin receptors (TfR), and receptors of ε-toxin produced by *C. perfringens* and α-toxin of *C.Septicum* (CPεTR and CSαTR, respectively) - through the development of new computational methods. By establishing a new analysis pipeline combining Bayesian inference, decision trees and clustering approaches, we indeed systematically classified single protein trajectories according to the type of confining energy landscape. This revealed the existence of only two distinct organization modalities: (A) confinement in a quadratic energy landscape for EGF, CPεT and CSαT receptors and (B) free diffusion in confinement domains resulting from the steric hindrance due to F-actin barriers for transferrin receptors.

The characterization of confinement energy landscapes by Bayesian inference furthermore revealed the role of interactions with the domain environment in cholesterol- and sphingolipid-rich domains with (in the case of EGFR) or without (for CPεT and CSαT receptors) parallel interactions with F-actin, to regulate the confinement energy depth. Strikingly, these two distinct mechanisms result in the same organization type (A). We furthermore revealed that the apparent domain sizes for these receptor trajectories resulted from Brownian exploration of the energy landscape in a steady-state like regime at a common effective temperature, independently of the underlying molecular mechanisms. These results highlight that the membrane organization in confinement domains may be more adequately described as interaction hotspots rather than rafts with abrupt domain boundaries.

Altogether, these results establish a new computational approach, which paves the way to the constitution of an atlas of energy landscape of membrane proteins and of their control mechanisms, and support a new general model for functional receptor confinement in membrane nanodomains.

## INTRODUCTION

The membrane nano- and micro-organization and its functional role is a longstanding riddle that attracts a considerable amount of research using increasingly sophisticated experimental and analysis tools (1–7). Among the most well-known confinement mechanisms in the membrane are i) the domains rich in cholesterol and saturated sphingolipids which provide the most favorable environment for numerous membrane proteins and receptors initially proposed by Brown and Rose (8) and Simons and van Meer (9) and dubbed lipid rafts, and ii) the domains defined by the steric hindrance provided by the underlying actin filaments which act as fences for membrane proteins with a protruding cytosolic part and by the membrane proteins attached to the actin cytoskeleton which act as pickets hindering the free diffusion of both proteins and lipids proposed by Kusumi and colleagues and designated as fences and pickets domains (10–13).

A functional role, in particular in signal transduction, was suggested for rafts(14–16). Although rafts were initially thought to be stable and a few hundred nanometers large, a series of publications reported the existence of small (a few tens of nanometers) and short-lived domains (2,17–20). The small size and transient nature of these domains lead to question whether the term raft was appropriate. Yet, in several cases, long-lived (> 1 s) and large (>100 nm) membrane domains are indeed observed (1). The connection between the domain content, cholesterol- and sphingolipid-rich, and their size is still unclear: in particular, whether the larger, long-lived domains arise from the coalescence of multiple small, short-lived ones, as suggested by Lingwood and Simons based on cross-linking experiments(16).

In addition, it was realized that the protein concentration in the membrane and in these domains in particular was much larger than initially thought which lead to describing these domains as protein-lipid composites and more frequently using the denomination raft or raft domain rather than lipid raft. Moreover, a number of additional membrane domains have been described such as caveolae, flotillin domains, clathrin-coated pits, protein assemblies, and tetraspanin-enriched domains which may also show cholesterol and/or sphingolipid dependence and some of them, *e.g*. caveolae or flotillin domains, may be considered as subcategories of raft domains, as reviewed in(6). In the following, we will use the generic term raft to designate all cholesterol- and sphingolipid-rich nanodomains.

Are these differences between small and large domain properties due to experimental artifacts, different mechanisms of membrane receptor confinement or variability between different cell types? Addressing this question is crucial, because this may influence the receptor activation process and the subsequent functional cell response involving critical signaling pathways, as is the case for the epidermal growth factor receptor EGFR(21–23). Although the organization of receptors into nanodomains has often been hypothesized as a general mechanism for regulating receptor activation, the role of molecular composition in the different types of confining domains is still not clear.

To improve our understanding of these domains we studied them from the perspective of the effective energy landscape seen by the proteins performing random walks within them. This should notably reveal if each given set of molecular composition and interactions leads to a specific energy landscape inside the domain or if the same type of energy landscape can be obtained through multiple composition and interaction pathways.

We implemented a Bayesian inference scheme to extract a diffusive profile and an effective interaction landscape as previously demonstrated in (24) from recordings of single-molecule trajectories of multiple membrane proteins. To ensure precise inference results, we performed the analysis of long receptor trajectories, typically 1000 points or longer using photostable nonblinking Eu-doped yttrium vanadate nanoparticles as labels (25). These long recordings ensure sufficient space sampling and exploration of the confinement borders -when relevant-by single receptors (26).

We have already shown that the confinement of ε-toxin and α-toxin receptors, CPεTR and CSαTR, in cholesterol- and sphingolipid-rich raft-type domains in MDCK cells could be approximated by an isotropic harmonic energy landscape with a stiffness value *k_r_* (25,27). The approach was then extended to infer large-scale energy and diffusivity landscapes at the total cell surface scale by exploiting the information stored in a large number of very short singlemolecule trajectories (28–30). The challenge in these inferences is that uncertainty in singlemolecule localization or density of displacements in space translates into non-easily predictable error on the diffusion and interaction landscapes. In our case, the high information content trajectories are provided by labeling of the receptors with bright and highly photostable Eu-doped yttrium vanadate Y0.6Eu0.4VO4 30-nm nanoparticles which yield a large number of trajectory points with a localization precision of about 30 nm and a temporal resolution of 50 ms (25).

We here aim at determining the effective interaction energy landscape experienced by membrane receptors inside different types of confining domains and relate its features to raft domain properties, to interaction with scaffolding proteins and with the cytoskeleton.

We focused on MDCK epithelial cells and implemented a quantitative approach based on Bayesian inference analysis of long trajectories to characterize the effective interaction energy landscape experienced by several membrane receptors confined inside domains resulting from different molecular mechanisms: epidermal growth factor receptor (EGFR) – a tyrosine kinase receptor with a single transmembrane domain, mostly present under a dimeric form at the cell membrane (31,32)) and possessing an actin-binding domain(33) -, CPεTR, CSαTR – the receptors of pore-forming *Clostridium perfringens* ε-(34) and *Clostridium septicum* α-toxins(35)-, and transferrin receptors (TfR)- a dimeric carrier protein for iron-binding transferrin (36). The interest in EGFR is particularly strong because of its role in proliferation and in numerous types of cancer (37) and is a common therapeutic target(38). The receptor of CPεT was found to be the hepatitis A virus cellular receptor 1 (HAVCR1)(39), which is also called T-cell immunoglobulin and mucin domain 1 (TIM-1) and is composed of an extracellular region consisting of an N-terminal immunoglobulin (Ig)-like domain attached to a mucin domain, a single transmembrane domain and a short C-terminal cytoplasmic tail(40). The CSαT has been shown to recognize glycosylphosphatidylinositol(GPI)-anchored proteins on the cell membrane(41)which are known to be found in rafts(20). While EGFR activity relies on its cholesterol-dependent clustering (42), and CPεTR and CSαTR are both present in cholesterol-rich domains (25), TfR has been reported to be associated with the cholesterol-poor membrane fraction (43).

Using a decision-tree and a clustering approach we showed that all three ε-, α-toxin, and EGF receptors experience a similar interaction landscape that can be described by a second-order polynomial, whereas transferrin receptors cluster in a different type of energy landscape better described by a fourth-order polynomial. Despite the fact that the molecular mechanisms involved for ε- and α-toxin receptors and for EGFR confinement are different – confinement in cholesterol- and sphingolipid-rich raft-type domains for ε- and α-toxin receptors and simultaneous confinement in raft-type domains and through binding to actin filaments for EGR receptors – both these confinement mechanisms create interaction landscapes with similar features.

Moreover, we showed that the description of confinement domains in terms of energy landscapes is particularly relevant. Indeed, we demonstrate that, in the case of second-order polynomial confinement, the experimentally observed domain size is (i) intimately related to the stiffness of the harmonic confinement energy experienced by the receptor and is (ii) an “apparent” domain size determined by the areas the receptor may explore given its thermal energy and the nanodomain energy landscape. This implies that the experimentally observed “apparent” domain size is not necessarily an actual physical feature of the domain reflecting an actual lipidic/proteic domain size, but is the result of the interactions giving rise to the confining interaction landscape and leading to the confinement of the receptor in the lower-energy areas of the domain compatible with its thermal energy.

## RESULTS

### Labeling and tracking EGF and transferrin receptors

We labeled individual Epidermal Growth Factor Receptors (EGFR) at the membrane of MDCK cells with a Y_0.6_Eu_0.4_VO_4_ nanoparticle-streptavidin (NP-SA) conjugate linked to biotinylated EGF ligands (Fig. 1A and Materials and Methods). This conjugate binds specifically to the EGFR extracellular domain and labeling conditions are adjusted so that only *ca*. 10 receptors per cell are effectively tagged (Supporting Information Fig. S1). We verified that the NPs labeling EGFR were exclusively present on the apical surface of the live MDCK cells by checking that they were localized in a plane a few μm above the coverslip surface (Fig. 1B). We tracked single NP-labeled EGFR and obtained uninterrupted trajectories of up to 240 s (*i.e*. 4600 points with an acquisition time *T_acq_* = 50 ms), due to the absence of blinking of these Eu-containing nanoparticle labels. The duration of the analyzed trajectories is in fact closer to 50 s because the combination of cell motions and mechanical instabilities of the microscope prevent the recording of spatially accurate trajectories for arbitrarily long durations, which leads to an exploration of the membrane with a typical density of 10,000 points/μm^2^ (Figure 1). The drift of the confinement domain (either due to mechanical instabilities or to nanodomain drifting inside the membrane) during long trajectories was removed by subtracting the average positions from the trajectories for each time interval. We only analyzed trajectory portions without measurable drift or after removing drift. A typical single receptor trajectory is shown in Fig. 1C. We observed that all tracked EGFR experienced a confined motion in an area with a typical radius *r*~250 nm, (Fig. 5), defined as the radius of a circle including 95% of the trajectory points.

**Figure 1.**
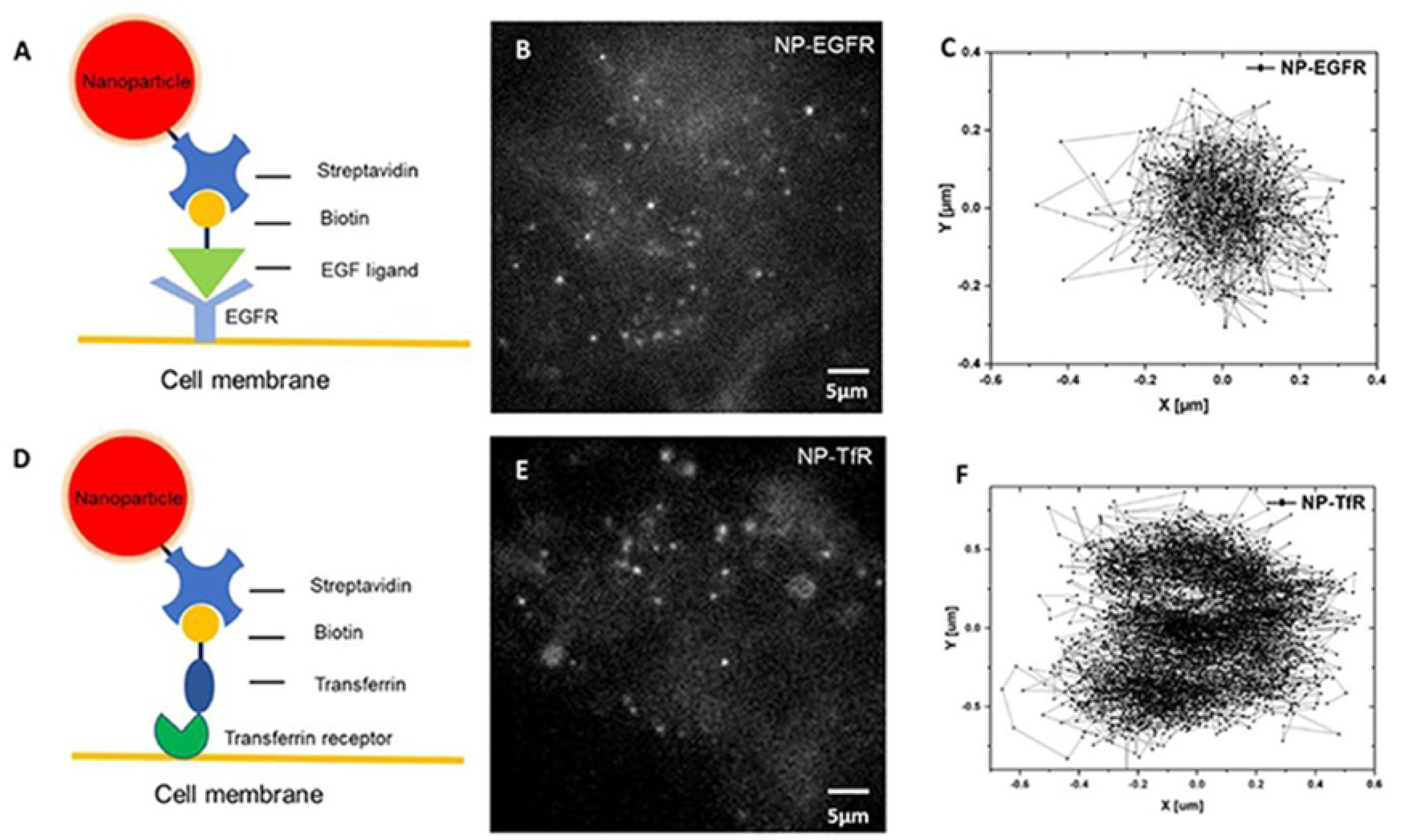
(A, D) Schematic of the Y_0.6_Eu_0.4_VO_4_ NP-streptavidin conjugate coupled to biotinylated EGF (A) and biotinylated transferrin (D) ligands binding to EGFR (A) and TfR (D), respectively, on the cell membrane. (B, E) A frame of the image sequence showing NPs labeling EGFR (B) and TfR (D), respectively, on the apical surface of live MDCK cells at 37 °C. (C, F) Trajectories of a single EGF receptor (C) and a single TfR receptor (F), respectively. Exposure time: 50 ms. Readout time: 1.3 ms.

We moreover labeled transferrin receptors (TfR) which, in contrast to EGFR, are known to reside outside of rafts(13,44) with NP-SA linked to biotinylated transferrin, and tracked TfR in the same way as EGFR, as shown in Fig. 1D-F. For all transferrin receptors (N=65), we observed hopping diffusion, *i.e*. short-lived confinement and frequent hopping from one confinement domain to the next (*e. g*. two hopping events and confinement in three different nanodomains are observed in a typical trajectory on Fig. 1F), in agreement with previous observations (13). This behavior has been attributed to “fences-and-pickets”-type confinement (11,12). The receptor motion is hindered by the fact that its intracellular domain bounces off the actin fences encountered and by its collisions with other proteins bound to filament actin. Frequent hops above the actin fences to adjacent confinement domains(11,13) show that the energy barrier to hop above the actin fences is low.

### Inference of the effective energy landscape

To quantitatively analyze these trajectories and measure the physical parameters controlling the receptor motions, we used the Bayesian Inference (BI) method presented by Masson et al.(24), which was applied to extract diffusivity and effective interaction energy landscapes of confined *Clostridium septicum* α-toxin receptor (CSαTR) and *Clostridium perfringens* ε-toxin receptor (CPεTR) trajectories in live MDCK cell membranes(25,27). Random walks within the membrane environment were modeled as a heterogeneous overdamped Langevin equation with spatially constant diffusivity and a conservative effective interaction force field. We thus reduce the complexity and heterogeneity of the membrane environment into diffusion coefficients and effective interaction energy landscapes. This method provides a simplified but effective description of the receptor dynamics within its confinement domain. In order to regularize the interaction field but also reduce the number of variables, the confinement effective interaction field was projected onto a polynomial basis. If there are no interactions, the interaction field will be constant. We have previously shown for simulated trajectories inside a flat interaction field with infinite barriers at the border that the Bayesian inference algorithm finds forces acting on the confined receptors that are close to zero, and vanishingly small values for the coefficients of the polynomial confinement potential(27).

The Bayesian inference implementation is described in detail in the Supporting Information and in our previous work (25,27). Briefly, the confinement domain is divided into subdomains of equal size, within which we assume the gradient of forces is negligible. The model being Markovian, the likelihood of the full trajectory is the product of the likelihood within each subdomain. Furthermore, the likelihood within each subdomain is directly the product of the likelihoods of all individual translocations within the subdomain. Finally, the likelihood of each translocation is the solution of the Fokker-Planck equation associated to the Langevin equation.

The diffusion coefficient was considered to be constant in all subdomains. Indeed, when the diffusion coefficient was allowed to vary between subdomains, the variations observed were not important(27). We initially determined the trajectory type by identifying the lowest possible order of the polynomial shape of the energy landscape sufficient to account for the experimental data. Rather than using Bayesian evidence to select the order of polynomials, we used the Kolmogorov-Smirnov (KS) test. This choice was motivated by the significant time required to properly sample the evidence of all models while the KS test demonstrated good performance (Supporting Information Fig. S2). For all the EGFR trajectories, there were no significant differences between fourth- and second order polynomials in the coefficients of the *x*^2^ and *y*^2^ terms, similarly to what we found in our previous work on CPεTR and CSαTR (25). We then defined the radial spring constant *k_r_* as *k_r_* = (*k_x_*^2^ + *k_y_*^2^)^1/2^ where (*k_x_, k_y_*) were inferred from the trajectories.

To go beyond this initial characterization, we here propose two procedures to systematically classify trajectories of different receptors: (i) a procedure based on a Bayesian inference decision tree which can distinguish between second- and fourth-order polynomials describing the effective interaction landscape and (ii) a clustering approach with no prior assumptions on the receptor motions. We show that clustering different receptors based on their trajectory properties and on the inferred energy landscape leading to these receptor motions can reliably discriminate between confinement types (“raft” versus “fences-and-pickets”) using statistics tools only and without requiring the use of pharmacological treatments as is currently done.

### Trajectory classification

#### Bayesian inference decision tree

We applied a Bayesian Inference Decision Tree (BIDT) approach previsouly demonstrated in (45) to attribute the motion observed in receptor trajectories to either free Brownian motion, confined motion in a harmonic polynomial or confined motion in a fourth-order polynomial landscape (Fig. 2). The method relies on a sequential approach to Bayesian testing allowing model identification through conditional tests (see Supporting Information). Türkcan and Masson have already demonstrated using simulated trajectories that the approach works for free and confined trajectories in harmonic and fourth-order energy landscapes(45). In the case of transferrin, most trajectories exhibit hopping. Thus, the trajectories need to be split into the successive confinement domains explored during the trajectory before the BIDT approach can be applied to each domain obtained after splitting. To show that the BIDT approach also works for trajectories involving hopping, we simulated a receptor moving in two adjacent domains with two harmonic energy landscapes on one hand and with two energy landscapes described by the equations below on the other (respectively Equation 1 and 2; Fig. 2A, 2B). The latter energy landscapes are flat in the center of the domain and exponential at the borders, as seen in Fig. 2B, we, therefore, call them flat-exponential or exponential energy landscapes.

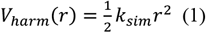

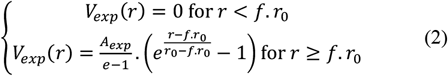

where *A_exp_* is the depth of the potential at *r*_0_ and is set at 4*k_B_T*, and *f* is the fraction of *r*_0_ where the flat potential starts to rise and is set at ⅓.

**Figure 2.**
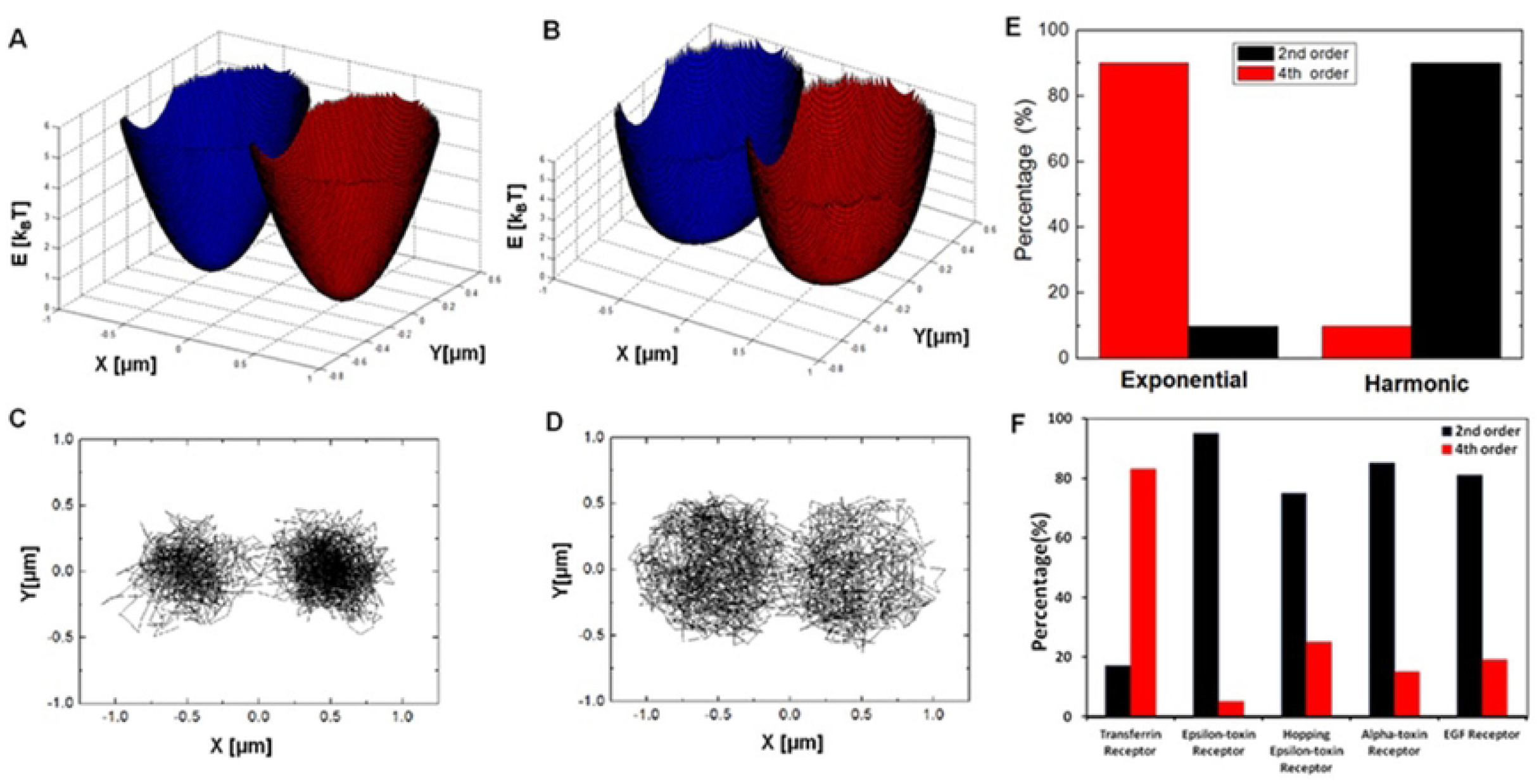
Adjacent confinement landscapes of the harmonic (A) and the flat-exponential type (B) used to simulate hopping trajectories (C) and (D), respectively, using N=3000, Δt=51.3 ms, D=0.1 μm^2^/s. (E) Decision-tree results for simulated trajectories involving two confining domains after splitting the trajectories in two parts using k-means. (F) Decision-tree results for experimental trajectories after splitting with k-means the trajectories involving multiple confinement areas.

We considered flat-exponential potentials in these simulations because this is the type of confinement landscape expected for a receptor moving inside a “fences-and-pickets”-like domain. Indeed, if the energy barriers to hop from one confinement domain to an adjacent one are defined by actin filaments underlying the membrane and by proteins bound to those filaments thereby hindering the receptor motion, it is reasonable to expect that for these receptors the energy landscape causing their confinement is flat in the center of the domain and rises abruptly at the domain borders.

50 trajectories of 3000 points of each kind were simulated, then split using k-means into 100 split trajectories for each condition. An example set of trajectories is shown in Figs. 2C and 2D below the respective energy landscapes. For these simulations, a diffusion coefficient of 0.1 μm^2^/s is used, typical of the one observed in the experimental trajectories (Fig. 4D). After splitting, the simulated trajectories are then fed into the BIDT algorithm. Figure 2E shows the results of the BIDT algorithm for the 100 trajectories that were obtained with harmonic potential energy profiles and the 100 trajectories obtained with exponential profiles. The trajectories that were initially produced using a harmonic profile are largely classified as being due to a second-order polynomial landscape, whereas the data produced with the flat-exponential confining profile are mostly identified as being due to a fourth-order rather than a second-order polynomial landscape. This confirms that a relatively flat energy landscape with abrupt energy barriers at its borders is more likely to be identified as a fourth-order polynomial one in our procedure and can be efficiently differentiated from confinement domains with a harmonic-like energy landscape. Consequently, this validates the BIDT procedure as an efficient trajectory classification method.

We then analyzed our experimental trajectories using the same BIDT approach. The splitting of the trajectories involving hopping was done using k-means with a domain number obtained by visual inspection of the trajectory. Note that, for experimental trajectories, it is crucial to first subtract any drift components that may be present, to avoid any improper classification. As shown in Fig. 3F, 65 transferrin, 40 ε-toxin and 20 α-toxin, 12 hopping ε-toxin and 21 EGF receptor trajectories were analyzed. The ε- and α-toxin data are the same as those analyzed in(25). We stress that the number of points, *N*, per trajectory used in this analysis was always above 500 and in most cases above 800. This ensures that the information amount contained in each trajectory is sufficient to determine the confinement landscape precisely enough so that the BIDT algorithm is able to treat more than 80% of trajectories correctly, according to our simulations.

**Figure 3.**
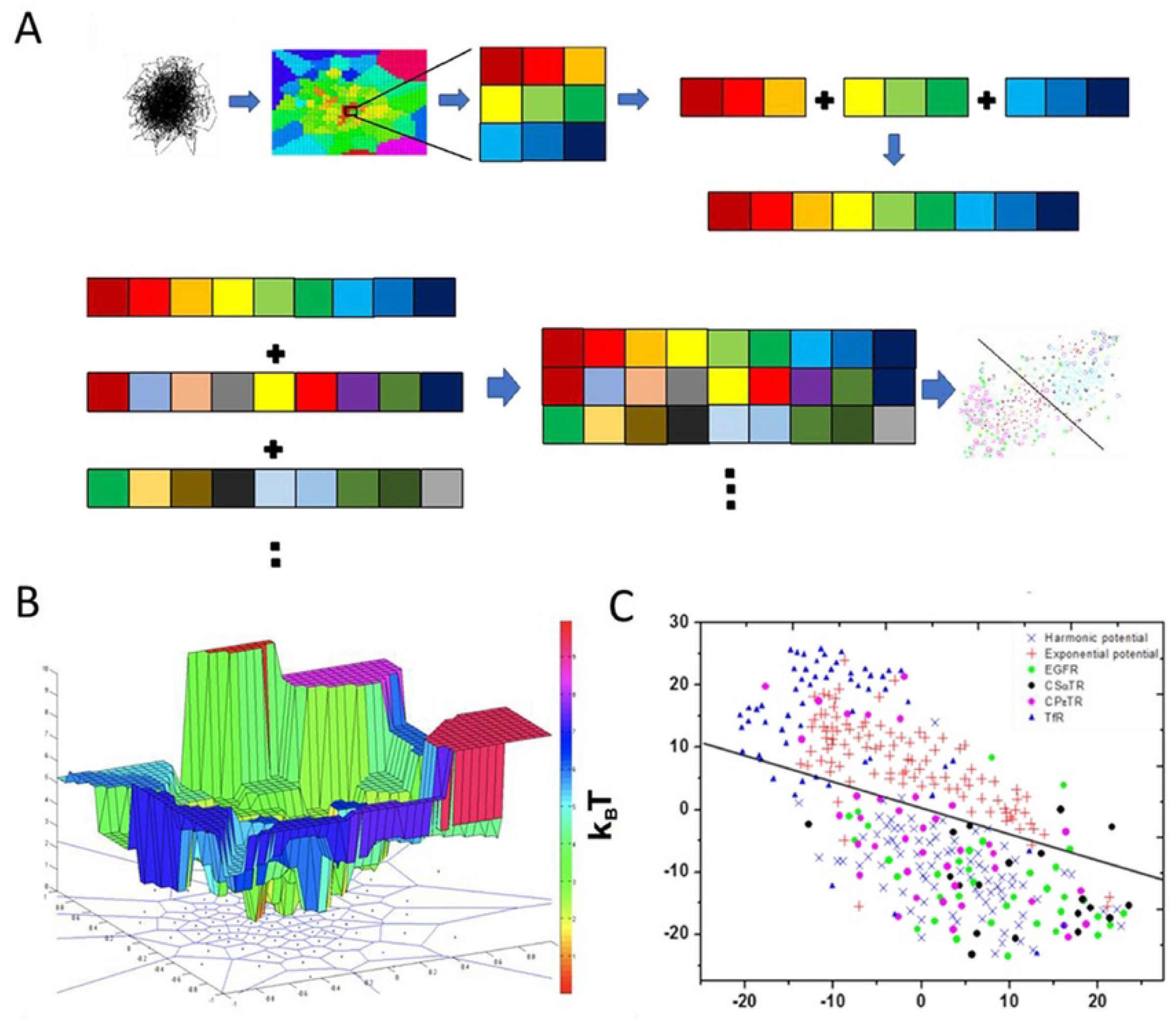
(A) Graphic representation of pre-treatment before t-SNE analysis(47). After Voronoi tessellation, the energy landscape map is inferred from the trajectory by Bayesian inference and projected on a mesh of 41×41 square subdomains. For reasons of clarity, only the analysis of the central part of the energy map is shown. In practice, each row of 41 energy values is put next to the previous one to create a single data vector containing 1681 values, which is used for PCA and then t-SNE analysis to obtain a 2D clustering. (B) Typical energy landscape map inferred from a trajectory with the subdomains obtained by Voronoi tessellation (blue line projection) and the square subdomains obtained after transformation into identical 41×41 square subdomains (colored mesh). (C) The t-SNE algorithm yields two clusters when applied to energy landscapes inferred from simulated trajectories with a harmonic confining landscape (blue crosses) and a flat-exponential landscape (red crosses). The solid line separates these two clusters. The potentials inferred from experimental trajectories of EGF, CSαT, and CPεT receptors are mainly projected onto the same cluster as that of the simulated data with a harmonic energy landscape, whereas trajectories of TfR receptors are projected on the same cluster as that of simulated data with flat landscape with exponential borders.

The results reveal a clear pattern. Raft-associated α-toxin and ε-toxin receptors are mostly attributed to harmonic energy landscapes (respectively 85%, 95%), as expected based on our previous results for these toxin receptors(25) This is also true for EGFR: 81% of the trajectories are identified as resulting from a harmonic energy landscape. Given that the ε-toxin and α-toxin receptors were determined to be confined in rafts in our previous work(25) and that EGFR are frequently found to be located in rafts in the literature(46), our results may hint that a raft-type nanodomain yields a second-order polynomial for the confinement energy landscape. The precise type of confinement of EGFR in MDCK cells was determined to be indeed raft-related as will be presented in the following section. In contrast, the raft-independent transferrin receptor trajectories are more likely to be identified as being due to a confining energy landscape described by a fourth-order polynomial (82%). These results imply that we can discriminate between raft (ε- and α-toxin receptors and EGFR, as demonstrated below) and non-raft associated receptors (transferrin receptors in our case) with a good -*i.e*. better than 80%-reliability degree.

#### Trajectory clustering

Although the analysis of single trajectories by the BIDT approach accurately distinguishes between raft-associated and non-raft confined receptors, it is not clear whether all the different raft-associated receptors are indistinguishable from each other based on their motions or can be separately classified depending on the receptor type. We thus propose a two-dimensional clustering approach to identify possible different confinement classes which may yield two (due to second- or fourth-order confining polynomials, as considered in the BIDT approach above) or more classes of receptor trajectories. We therefore implement the t-distributed stochastic neighbor embedding (t-SNE) clustering tool(47) to the clustering of confining energy landscapes extracted from single-molecule trajectories.

In the following, we used a more elaborate Bayesian Inference approach (see Materials and Methods), using a Voronoi division of the explored nanodomains (Fig. 3A) in such a way that each Voronoi subdomain contains an equal number of trajectory points, to obtain a mapping of the energy landscape and of the diffusivity which is here allowed to vary in each subdomain. This yields diffusivity and energy landscape maps (Fig. 3B) which were then transformed into 2D 41×41 square matrices of identical dimensions. These 2D energy landscapes were transformed into 1681-long 1D vectors as shown in Fig. 3A and were subsequently sorted in clusters in a two-dimensional space through principal component analysis (PCA) followed by a t-distributed stochastic neighbor embedding (t-SNE) algorithm(47), which classifies trajectories on the basis of the energy landscape shape and diffusivity (Materials and Methods).

We validated this approach by first simulating trajectories of both types (receptor experiencing confinement in a harmonic landscape and in a flat landscape with exponential boundaries) with the same constant diffusivity of 0.1 μm^2^/s and determining their position in the 2D plane (Fig. 3C). This shows that these two types of simulated trajectories form two different clusters with negligible overlap. Based on these simulations, we then defined a line separating these two clusters, into which each experimental trajectory is affected as a function of its position in the clustering plane relative to this line. We could then determine the percentage of the trajectories of a given type of receptors located inside each of these two clusters.

The clustering of the experimental data demonstrates that EGF receptors, as well as α-, and ε-toxin receptors have a different behavior than transferrin receptors, as expected based on the BIDT results. Indeed, 92% of EGFR, 100% of α-, and 95% of ε-toxin receptor trajectories fall inside the cluster of harmonic energy landscapes, while 82% of transferrin receptor trajectories fall inside the flat-exponential cluster (Fig. 3C). Interestingly, no relevant sub-clustering can be made to separate EGFR, α-, and ε-toxin receptors (Fig. 3C). This indicates that EGFR and α- and ε-toxin receptors are confined in harmonic-like energy landscapes, which may consequently imply that EGFRs and toxin receptors may reside in domains with similar dynamics properties, even though the underlying set of interactions may possibly differ.

Our previous work has shown that this clustering approach can also be applied to 3362-long vectors including both the confinement energy map and the diffusivity map (data not shown). This yields very similar results demonstrating that the main feature differentiating the trajectories is the confining energy landscape and not the diffusion properties.

Given the BIDT classification and the clustering results, EGFR and α- and ε-toxin receptors seem to share a common organization in MDCK cells. As we have previously shown with experiments under cholesterol oxidase and sphingomyelinase incubation, α- and ε-toxin receptors are located inside cholesterol- and sphingolipid-rich rafts(25). The molecular origin of EGFR confinement needs therefore to be determined to be able to draw conclusions about whether EGFR and α- and ε-toxin receptors share the same domain type, *i.e*. rafts. We thus assayed quantitatively the EGFR domain properties to decipher the confinement mechanisms leading to the same type of energy landscape.

##### Confinement of EGF receptors is raft- and actin meshwork-dependent

After identifying that EGFR are mainly confined in a harmonic energy landscape, we determined the stiffness of this potential, *i.e*. the spring constant *k_r_*, and the EGFR diffusion coefficient *D_Inf_* by Bayesian inference, as discussed in the section Inference of the effective energy landscape and in (25). The confinement domain surface *A* is also determined independently as discussed above. We then compared these measurements with raft-associated receptors CSαTR, CPεTR and non-raft transferrin receptors (TfR).

To investigate the molecular origin of this potential, we first disrupted cholesterol- and sphingomyelin-rich raft domains by cholesterol oxidase (ChOx) and sphingomyelinase (SMase) and then depolymerized the actin cytoskeleton by latrunculin B (Lat B). When the membrane was cholesterol depleted, we observed a significant reduction of confinement (Fig. 4A) and a significant increase of domain area (Fig. 4C) of EGF receptors. After 30 min of treatment, the average spring constant < *k_r–EGFR_* > is reduced by 83±4% (< *k_r–EGFR_* >=0.67±0.08 pN/μm, *N*=21 without, and 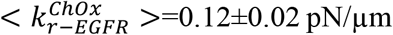, *N*=20 with ChOx incubation), and the average diffusion coefficient < *D_inf_* > increased from 0.065±0.006 μm^2^/s to 0.215±0.019μm^2^/s (Fig. 4D). Correspondingly, the average domain area < *A* > increased from 0.26±0.02 μm^2^ to 1.46±0.18 μm^2^ (Fig. 4C). These results demonstrate the central role of cholesterol in the origin of the confinement effect. To test if the EGFR confining domains are cholesterol- and sphingomyelin-rich raft domains, we used sphingomyelinase incubation to disrupt them. While sphingomyelinase treatment had a similar effect to cholesterol oxidase on CSαTR and CPεTR dynamics(25), in the case of EGFR, it results in a drastic reduction of EGF receptors present in the membrane, as revealed by the lack of visible EGFR after our labeling protocol (Fig. S3). This points to a key role of sphingomyelin, which seems essential for the EGFR insertion in nanodomains and/or its addressing and stabilization at the membrane.

**Figure 4.**
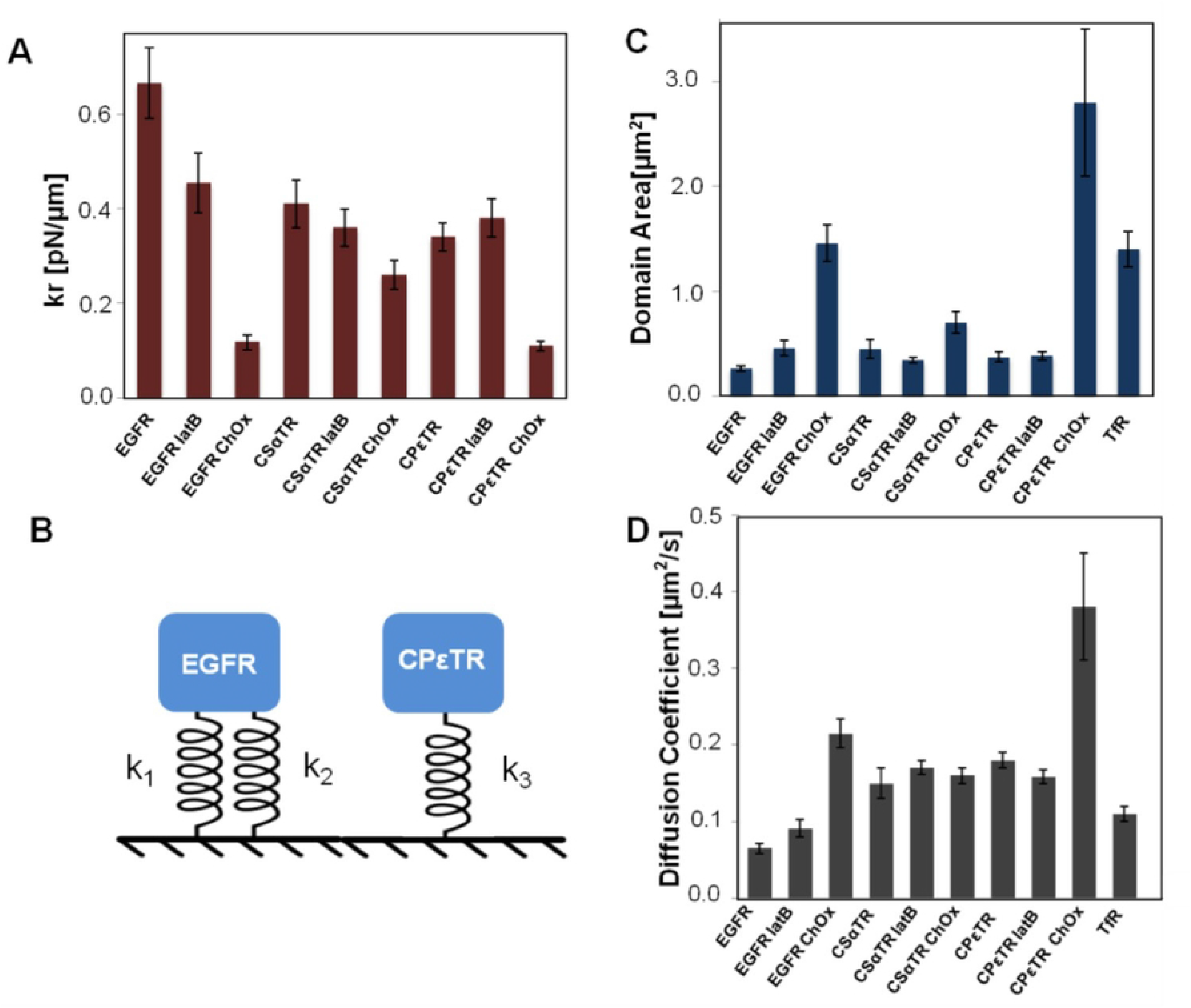
Effect of cholesterol extraction and actin depolymerization on the motion of EGF, CSαT, CPεT and Tf receptors. Cells were treated with 5μM latrunculin B and 20U/ml cholesterol oxidase, respectively. (A) Spring constants inferred using Bayesian inference as explained in the text. (B) The EGF receptor confinement can be modeled with two parallel springs, the stiffness of each spring, k_1_ and k_2_, being related to the stiffness of the actin meshwork and to the stiffness resulting from the confining energy landscape of the raft domain, respectively. The total stiffness, k_r_, is then-equal to the sum of k_1_ and k_2_. The CSαT and CPεT receptor confined can be described with a single spring system. (C) Domain area. (D) Diffusion coefficient.

After depolymerizing the actin meshwork, the average < *k_r–EGFR_* > is reduced by 34±11% 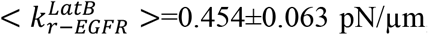, *N*=20), the average < *D_inf_* > increased to 0.091±0.011 μm^2^/s, and the average < *A* > increased to 0.456±0.068 μm^2^. We can thus deduce that the stiffness of the effective confining landscape in which EGF receptors move also includes a part of cytoskeleton confinement. We attribute this to direct EGFR-actin cytoskeleton interaction through the actin-binding domain of EGFR(33).

Based on these observations we can thus model the EGFR confinement with a parallel two-spring system as shown in Fig. 4B, where the effective elasticity due to the confining energy landscape *k_r_* = *k*_1_ + *k*_2_ results from two components *k*_1_ and *k*_2_, describing respectively the interaction with raft domains and F-actin. In contrast, the depolymerization of the actin meshwork results in no significant change in the case of CSαTR and CPεTR (25). In this case, the confinement results purely from receptor/raft interactions, as discussed in Ref. (25) that can be effectively described by a single spring model (Fig. 4B).

##### Confinement modeling of EGF, CSαT, CPεT receptors

The nature of the link between the confinement and the energy landscape is not clear: receptors could either be trapped in a nanodomain with boundaries, in which they experience a spring-like energy landscape or the domain could be the observable result of the receptor/lipid/protein interactions creating the confining energy landscape. In the first case, the confinement domain size would reflect the raft spatial ultra-structure, while in the second case it would only be due to the strength and the range of the confining interactions with the raft domain itself and/or with the underlying cytoskeleton meshwork.

As observed in Fig. 1C, most of the trajectory areas of raft-confined EGF receptors do not display any preferential direction and the confinement domain is approximately a circle, whose center can be determined by averaging the positions of all points. The potentials determined by BI are furthermore isotropic (*k_x_* ≈ *k_y_*) and the resulting potential energy is thus:

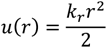

If we assume that the trajectories are long enough to reflect a situation of thermodynamic equilibrium at an effective temperature *T_eff_*, we can derive the probability density of the position of a receptor as a function of r and of the spring constant *k_r_*:

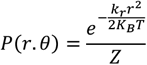

with *Z* the partition function defined to ensure that:

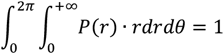

In our experiment, we defined the confinement area through the radius of a circle containing 95% of the total trajectory points. We thus have:

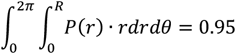

which yields the following scaling law:

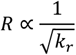

Consequently, if the confinement results purely from the interactions creating the spring-like potential and is at steady state, we expect that the radius of the confinement domain will be inversely proportional to the square root of the spring constant.

The comparison of the experimental values of *R* and the spring constant value *k_r_* obtained by BI for each trajectory are presented in Fig. 5 for EGFR, CSαTR and CPεTR before and after inhibition by ChOx and latrunculin B and reveal a good agreement with this prediction. This indicates that the organization of these receptors can indeed be described as a steady state process and reveals that the two parameters *k_r_* and *A* are correlated and both describe the same phenomenon: the confinement area *A* is consequently not determined by any physical boundaries of the nanodomains. Strikingly, this observation is a shared feature for different receptors and in different conditions, *i.e*. with or without partial actin meshwork or raft disruption, which leads to hypothesize that this confinement organization may be a universal feature in cells, independent of the underlying molecular events causing the confinement.

**Figure 5.**
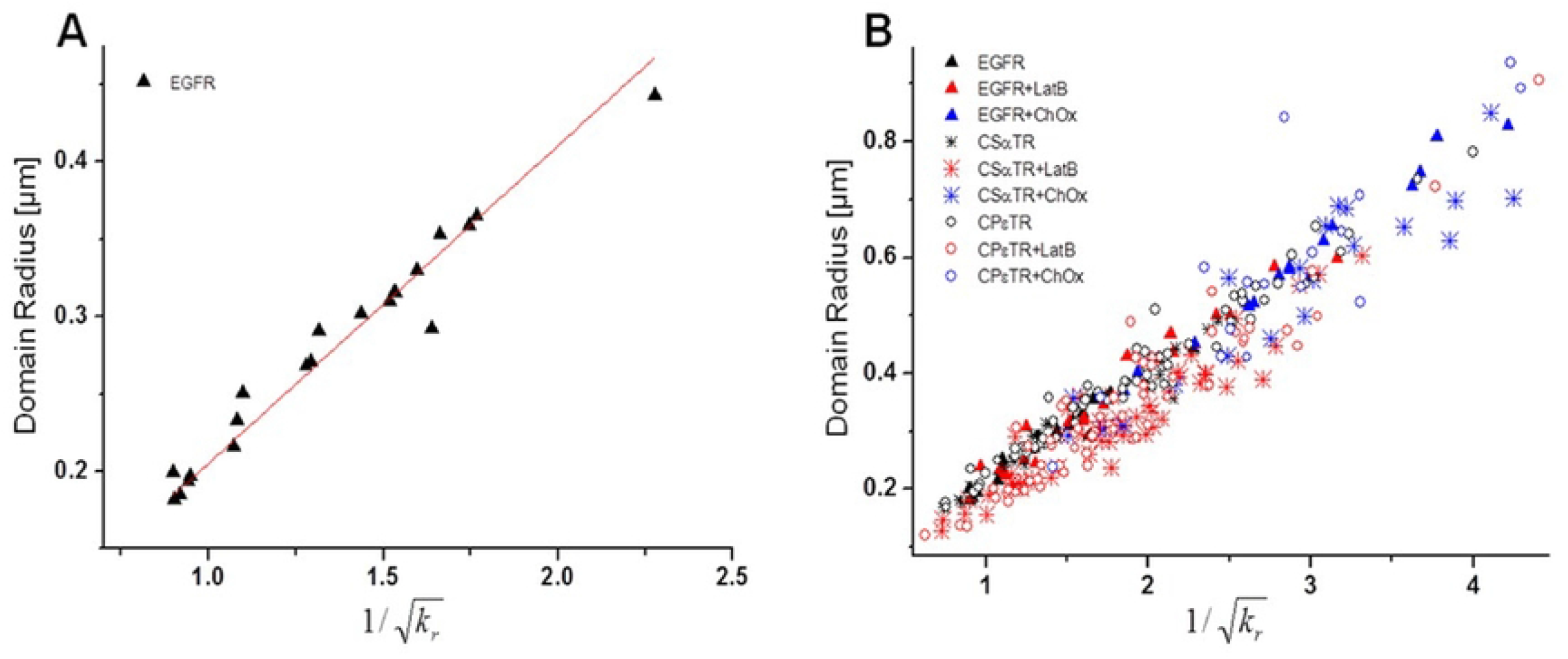
The domain radius is proportional to the reciprocal square root of the spring constant of the confining energy landscape. (A) EGFR. (B) All three receptor types, EGFR, CSαTR and CPεTR, respect this relation between domain radius and stiffness of the energy landscape even after disruption of rafts by cholesterol oxidase (ChOx) and depolymerization of the actin cytoskeleton by latrunculin B (Lat B).

## DISCUSSION

We identified through single-molecule tracking distinct molecular confinement mechanisms for different membrane receptors. We notably demonstrated unambiguously through Bayesian inference, that EGF, CPεT, and CPαT receptors were confined by similar quadratic energy landscapes. Moreover, we quantitatively measured the stiffness of these energy landscapes and revealed the multiple molecular origins of their shape in the case of EGFR. The latrunculin B treatment indeed reduced the stiffness of the energy landscape experienced by EGFR – indicating the F-actin implication - to a value close to the stiffness observed for the two toxin receptors which did not show any dependence on F-actin (Fig. 4A and (25)). This indicates that these 3 receptor types may be located in the same, cholesterol-sphingomyelin-rich domains.

Our results lead us to identify three types of membrane receptor nano-organization (Fig. 6): (i) the confinement in cholesterol-sphingolipid-rich raft nanodomains (for the two toxin receptors studied here), (ii) the simultaneous confinement in cholesterol-sphingolipid-rich nanodomains and attachment to cytosolic proteins below the membrane, such as F-actin (for EGFR), (iii) semi-free “hop” diffusion outside rafts and motion limited by actin-related barriers only observed for transferrin receptors. Although these two latter types of organization are cytoskeleton-dependent, they have significantly different functional consequences: EGFR are associated to nanodomains submitted to a strong interaction with the cytoskeleton, while transferrin receptors diffuse semi-freely in the membrane, only limited by actin barriers, according to the “fences-and-pickets” model.

**Figure 6.**
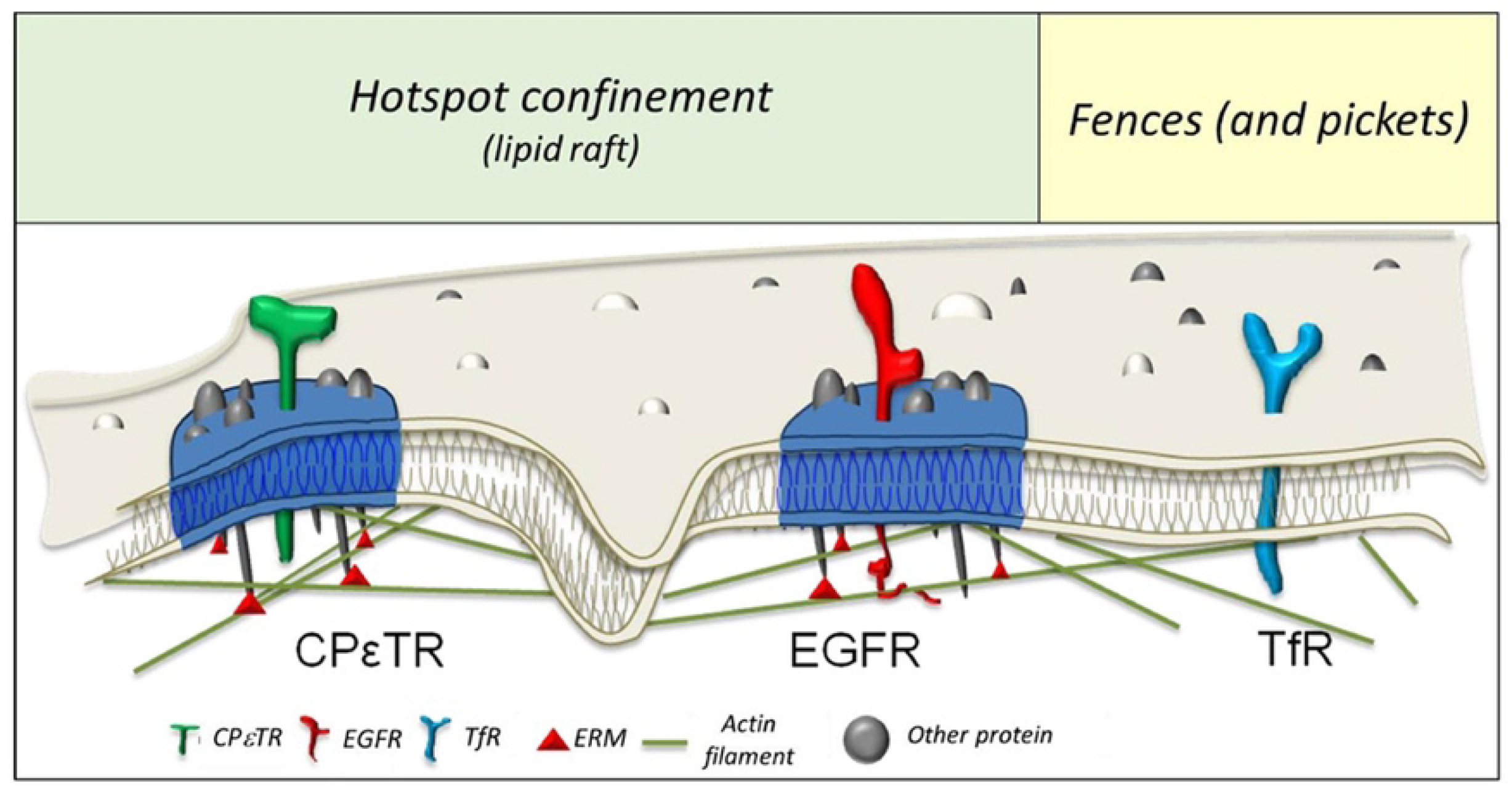
Three molecular models of membrane protein organization: (i) confinement in cholesterol- and sphingolipid-rich rafts (left, CPεT and CPαT toxin receptors), (ii) confinement both due to rafts and F-actin meshwork (center, EGFR) and (iii) diffusion and hoping over Factin dependent barriers (right, transferrin receptors) resulting in two functional organization types: quadratic energy landscape confinement in hotspots in cases (i) and (ii) and hop diffusion in a flat energy landscape with abrupt barriers (fences and pickets model) in case (iii). ERM stands for proteins of the Ezrin-Radixin-Moesin family, which mediate interactions between the plasma membrane and the actin cytoskeleton(55)

Although EGFRs have been reported in other systems to be present in different membrane nanodomain types (48–50), we here observed only one confinement type, indistinguishable from the CPεTR and CPαTR confinement in cholesterol-sphingomyelin-rich nanodomains in terms of the shape of the confining energy landscape. This observation, together with the cholesterol and sphingomyelin depletion data, indicates that, in our experimental conditions and cell type, EGFR receptors are present in cholesterol-sphingomyelin-rich domains. Note that, in our case, the receptor labeling is achieved through binding of the EGFR ligand EGF to the receptor, *i. e*. we are observing the motion of ligand-bound EGF receptors. Although these receptors are not necessarily activated receptors - two EGF molecules are required to activate a receptor dimer-, we cannot exclude that the motion and confinement characteristics of non-ligand-bound receptors may be different.

Interestingly, although EGFR and the two toxin receptors are confined through different mechanisms at the molecular level, they share similar confinement energy landscapes, which may have comparable functional roles. These results represent the first step in constituting an atlas of membrane receptor organization at the molecular scale, for which we studied 3 distinct mechanisms. However, at the functional scale, these mechanisms result in only 2 different organization types in which all examined receptors could be classified: type A with a confinement in a quadratic energy landscape at an effective temperature *T_eff_* relating the energy landscape stiffness to the domain radius, and type B with a fences-and-pickets like organization described by a confining energy landscape that is flat in the center and increases abruptly at the border. Indeed, non-raft receptors are classified as being submitted to a 4^rth^ order rather than a 2^nd^ order energy landscape by the BIDT approach (Fig. 2) and they cluster in a separate cluster from the raft-associated receptors which are all found in the same cluster (Fig. 3). This points to the ability of our method to identify the membrane localization of receptors in living cells – inside nano-organization domains based on molecular interactions versus domains defined by steric hindrance effects -, based on their motion only, without the further requirement of multiple pharmacological treatments like cholesterol oxidase and sphingomyelinase to be able to determine the raft or other nature of the confinement.

We here discuss the energy landscape experienced by confined receptors without *a priori* taking into account possible raft subcategories. This however does not mean that all three raft-confined receptors we examined are found in the same raft domains. Yet, if they are located in different raft subcategories, they are here found - within the limits of our experimental precision and subsequent trajectory analysis - to experience the same energy landscape.

All raft nanodomain-associated receptors of our data set thus share common organizational properties, but it should be stressed that other types of molecular-interaction-related confinement (*e.g*. rafts versus tetraspanin-enriched domains(51) or protein molecular assemblies(52) could possibly lead to alternative functional confinement types.

For the type A organization, we established a scaling law between the nanodomain radius and the potential stiffness, independent of the nature of the confined receptor (Fig. 5). This behavior is the signature that the organization of the membrane receptors can be viewed as having reached an apparent steady state, implying the existence of an effective temperature *T_eff_*. This notably indicates that their average distribution at the membrane is determined only by the energy landscape resulting from the presence of cholesterol-sphingolipid-enriched nanodomains and - in the case of EGFR - from binding to the actin meshwork. This generalized scaling law behavior is respected in various experimental conditions, in the presence of cholesterol oxidase, sphingomyelinase, and latrunculin B, as well as for three different types of receptors. Furthermore, *T_eff_*, though different from the actual temperature *T*, is unchanged (Fig. 4) in all tested situations: different receptors, pharmacological treatments. As mentioned in Ref. (53), the existence of active energy-consuming processes in solution can lead to an effective temperature *T_eff_ > T*, which controls the motion of passively diffusing particles. This may thus indicate the existence of active mechanisms in eukaryotic cells regulating effective membrane hydrodynamic properties and consequently membrane receptor mobility and thus leading to the formation of similar functional domains, or hotspots, for different unrelated membrane proteins. Strikingly, these observations remain true for different classes of confinement molecular mechanisms, cytoskeleton dependent or not, leading to common organization properties, which are thus mostly independent of the underlying interactions, and may indicate the existence of common functional features, such as the regulation of activity by nano-organization modulation, as hypothesized in the case of EGFR(50).

The question of the size of membrane micro- and nanodomains and in particular of raft domains has been controversial for a long time(18). The relation between the confinement stiffness *k_r_* and the domain radius we revealed (Fig. 5), highlights the fact that this question may not be relevant. Our data show that the observed domain size can be seen as the extension of the exploration of the membrane nanodomain by the receptor submitted to a quadratic energy landscape: therefore, the domain size does not directly reflect the physical size of cholesterol-sphingolipid-enriched membrane domains, but how far the receptor can move away from the domain center into higher energies inside the energy landscape given the thermal kinetic energy is possesses at the membrane effective temperature *T_eff_*. Indeed, the stiffer the confining energy landscape as a function of location, the more improbable it is for the receptor to reach locations far from the domain center and the smaller the domain radius will be. We can therefore consider the observed confined trajectory radius as an apparent domain radius. Indeed, the observed or apparent confinement radius (200-500 nm) indicates the stiffness of the energy landscape which reflects the strength of the receptor/lipid/protein/cytoskeleton interactions. We may thus conclude that the stiffness of the energy landscape is a more relevant parameter than the domain size characterizing both the confining domains and the distribution of proteins at the cell membrane.

Moreover, our results are consistent with the absence of organized raft nanodomains with abrupt frontiers: a locally varying mixture of different lipid types and proteins could result in such confinement, only determined by the characteristics of the interactions between the receptors and the lipidic-proteic membrane environment. This sheds new light on the concept of cholesterol-sphingolipid-rich domains (rafts) and might conciliate previous apparently contradictory findings from the literature. In contrast with the common picture of nanodomains, notably those observed in artificial lipid vesicles (54), we here demonstrate that their existence does not imply the presence of abrupt edges, separating two different lipid phases. The term “raft” also suggests the existence of nanodomains with abrupt barriers acting as functional signaling platforms (16) and may seem to be misleading. These domains should rather be seen as hotspots of sphingolipid, cholesterol and protein/receptor enrichment, which mediate the confinement properties.

Altogether, these results point to a renewed model for receptor organization in cholesterol-sphingomyelin-rich confinement nanodomains: the receptor motion in these domains may simply be described by free receptor diffusion inside a quadratic energy landscape with receptor localization resulting from the Boltzmann distribution inside this energy landscape. We can thus propose an explanation for the reported EGFR translocation between different domain types (from sphingolipid-rich to caveolar domains) after different stimulations (50): the stimulus may induce a modification of the effective stiffness of different interaction hotspots, by regulating the interactions between their components, leading to a transition towards another steady state in a new energy landscape.

## CONCLUSIONS

Altogether, these results support a new general model of the nano-organization of membrane proteins, such as EGFR, CPεT or CPαT receptors, in functional hotspots: though possibly created through distinct molecular mechanisms, these interaction hotspots may lead to similar organizational properties, determined by an equilibrium distribution in quadratic effective energy landscapes. Our approach, relying on innovative quantitative tools of single-molecule trajectory analysis, could thus constitute the basis of an atlas of membrane proteins, based not only on identified sets of interaction, but on their physiologically relevant organization and confining energy landscape at the membrane. This may contribute to the further understanding of the functional role of membrane organization in normal or pathological contexts.

## MATERIALS AND METHODS

### Nanoparticle preparation

Y_0.6_Eu_0.4_VO_4_ nanoparticles were prepared, functionalized, and coupled to proteins, as in our previous work (25,26). The nanoparticles are ellipsoidal with typical sizes of 30 nm. As described in (25), we coupled aminopropyltriethoxysilane (APTES)- coated europium-doped nanoparticles to ε-prototoxin produced by *C. perfringens* bacteria (CPεT), α-toxin produced by *C. septicum* bacteria (CPαT), or streptavidin, via the amine reactive cross-linker Bis(sulfosuccinimidyl)suberate (BS3). In a first step, APTES-coated nanoparticles reacted with BS3 in large excess to avoid crosslinking between nanoparticles. In a second step, the BS3-conjugated nanoparticles (NP) reacted with amine groups on the target proteins at a chosen coupling ratio (NP:ε-prototoxin 1:3, NP:α-toxin 1:5, NP:streptavidin 1:11). This second step is performed rapidly to avoid hydrolysis of the BS3 NHS active groups. To obtain EGF-NP and transferrin-NP conjugates, we further incubate NP-streptavidin complex with biotinylated EGF (Thermofisher) or biotinylated transferrin (Sigma-Aldrich) at 37°C with a molar ratio of 1:3 for 1h. We then remove unbound biotinylated ligands by high speed centrifugation for 80 min at 16,000 g and retain the pellet which is redispersed in phosphate buffer saline pH 7.4.

### Cell culture, labelling and pharmacological treatments

We cultured Madin Darby Canine Kidney (MDCK) cells in Dulbecco’s Modified Eagle’s medium (DMEM) with 10% fetal bovine serum and 1% penicillin-streptomycin at 37°C and 5% CO_2_. Cells were then transferred onto glass coverslips and starved overnight in FBS-free culture medium. The culture medium was then replaced immediately before imaging by an observation medium (OM: Hank’s Balanced Salt Solution (HBSS) + 10 mM HEPES) to avoid fluorescence due to the culture medium.

Cells were then incubated with 0.04 μM NP-labeled EGF, respectively Tf, for 15 min at 37°C, and then rinsed three times with observation medium to remove non-bound nanoparticles. In these conditions, the concentration of visible receptor-bound nanoparticles at the apical cell membrane was low enough to observe single NPs (typically 10 NPs per cell, Figure S1).

Pharmacologically treated cells were respectively incubated in OM containing 500 nM latrunculin B (Calbiochem, Millipore, Billerica, MA), 20 U/mL cholesterol oxidase (Calbiochem, Millipore), or 10 U/mL sphingomyelinase (Calbiochem, Millipore) for 30 min at 37°C before labeling with the nanoparticles and starting the imaging.

### Single receptor imaging and trajectory recording

Tracking experiments were performed with a wide-field inverted microscope Zeiss Axiovert 100 (Zeiss Oberkochen, Germany), or Olympus IX-81 inverted microscope (Olympus America, Center Valley PA, USA) equipped with a 63×, NA=1.4 oil immersion objective and an electron-multiplying charge-coupled device (Quant EM:512SC; ROPER Scientific, Trenton, NJ). The Y_0.6_Eu_0.4_VO_4_ nanoparticles are excited with an Ar^+^ ion laser using the 465.8 nm line and their emission is collected through a 617/8 filter (Chroma Technology, Bellows Falls, VT). We used white-light transmission imaging to focus on the cells, then switched to fluorescence imaging and searched upwards for the focal plane where NPs are in focus. The nanoparticle emission signal was only observed at a single focal plane, at the apical membrane of the live MDCK cells. Images were recorded at a 19.5 Hz frame rate (exposure time: 50 ms; read out time: 1.3 ms) with an excitation intensity of 0.25 kW/cm^2^ at 37°C.

In a low concentration regime, (~10^−3^-10^−2^ NP/μm^2^, Figure S1), the receptor positions in each frame were determined from a 2D Gaussian fit to the diffraction pattern of the nanoparticles with an algorithm run in a custom MATLAB routine (Mathworks, Natick, MA), yielding a typical 30 nm localization accuracy. Single trajectories were then unambiguously reconstructed based on a closest neighbor criterion, since, in a low concentration regime, no nanoparticle crossing can be observed for any labeling. The mean displacement, possibly due to mechanical drift, was measured for each trajectory, then subtracted before any analysis.

### Trajectory clustering

We first use a Voronoi-based division of the nanodomain into subdomains. First, the trajectories were clustered through a k-means algorithm(56), which provides seed points for the Voronoi tessellation. In this case, the subdomain size depends on the density of trajectory points in each location inside the nanodomain. We then inferred by Bayesian inference (see Supporting Information) the potential energy value in each subdomain. In this case, the diffusion coefficient is free to vary in each subdomain. This ensures a homogeneous number of trajectory points per subdomain which implies inferring the potential energy value in each subdomain with similar uncertainty. This yields, for each trajectory, potential energy points for each Voronoi subdomain (Fig. 3B). To be able to compare the potential energy maps with each other, these Voronoi potential energy maps are all transformed into 2D 41×41 square subdomain matrices (Fig. 3A,B). This yields subdomain sizes that, when superimposed onto the Voronoi division of the potential energy, are approximately half the size of the smallest typical Voronoi domain. This is small enough to conserve the heterogeneity of the data obtained from different trajectories, and large enough to avoid oversampling the inferred potential energy. The 41×41 subdomains give us a 1681-dimensional data set for the potential map and an equivalent data set for the diffusivity map, hence an overall 3362-dimensional data set per trajectory (Fig. 3A). This 3362-dimensonal data is reduced to 30 dimensions via a principle component analysis (PCA), to retain the highest degree of heterogeneity between data points before further analysis.

We then use a t-distributed stochastic neighbor embedding (t-SNE) algorithm to map the heterogeneity of our 30-dimensional data set onto two dimensions to sort individual trajectories in clusters reflecting the type of the energy landscape that led to the observed motions on a statistically reliable basis(47). The data points representing the individual trajectories were then plotted on a two-dimensional surface so that the distances between a point and all the other points reflect the similarities between the points quantified by a Student’s t-distribution.

## ACKNOWLEDGEMENTS

We thank R. Mohammedi and T. Gacoin (Laboratoire de Physique de la Matière Condensée, Ecole Polytechnique, CNRS UMR7643) for synthesizing and providing nanoparticles used in this study.

## References

1. Carquin M, D’Auria L, Pollet H, Bongarzone ER, Tyteca D. Recent progress on lipid lateral heterogeneity in plasma membranes: From rafts to submicrometric domains. Prog Lipid Res. avr 2016;62:1–24.

2. Eggeling C, Ringemann C, Medda R, Schwarzmann G, Sandhoff K, Polyakova S, et al. Direct observation of the nanoscale dynamics of membrane lipids in a living cell. Nature. 26 févr 2009;457(7233):1159–62.

3. Honigmann A, Pralle A. Compartmentalization of the Cell Membrane. J Mol Biol. déc 2016;428(24):4739–48.

4. Krapf D. Compartmentalization of the plasma membrane. Curr Opin Cell Biol. août 2018;53:15–21.

5. Levental I, Veatch SL. The Continuing Mystery of Lipid Rafts. J Mol Biol. déc 2016;428(24):4749–64.

6. Lu SM, Fairn GD. Mesoscale organization of domains in the plasma membrane – beyond the lipid raft. Crit Rev Biochem Mol Biol. 4 mars 2018;53(2):192–207.

7. Kalappurakkal JM, Sil P, Mayor S. Toward a new picture of the living plasma membrane. Protein Sci. juin 2020;29(6):1355–65.

8. Brown D, Rose J. Sorting of GPI-Anchored Proteins to Glycolipid-Enriched Membrane Subdomains during Transport to the Apical Cell Surface. Cell. 68^e^ éd. 1992;533.

9. Simons K, Van Meer G. Lipid sorting in epithelial cells. Biochemistry. 23 août 1988;27(17):6197–202.

10. Fujiwara T, Ritchie K, Murakoshi H, Jacobson K, Kusumi A. Phospholipids undergo hop diffusion in compartmentalized cell membrane. J Cell Biol. 10 juin 2002;157(6):1071–82.

11. Fujiwara TK, Iwasawa K, Kalay Z, Tsunoyama TA, Watanabe Y, Umemura YM, et al. Confined diffusion of transmembrane proteins and lipids induced by the same actin meshwork lining the plasma membrane. Bassereau P, éditeur. Mol Biol Cell. avr 2016;27(7):1101–19.

12. Kusumi A, Nakada C, Ritchie K, Murase K, Suzuki K, Murakoshi H, et al. Paradigm Shift of the Plasma Membrane Concept from the Two-Dimensional Continuum Fluid to the Partitioned Fluid: High-Speed Single-Molecule Tracking of Membrane Molecules. Annu Rev Biophys Biomol Struct. 1 juin 2005;34(1):351–78.

13. Sako Y, Kusumi A. Compartmentalized structure of the plasma membrane for receptor movements as revealed by a nanometer-level motion analysis. J Cell Biol. 15 juin 1994;125(6):1251–64.

14. Simons K, Ikonen E. Functional rafts in cell membranes. 1997;387:4.

15. Simons K, Toomre D. Lipid rafts and signal transduction. Nat Rev Mol Cell Biol. oct 2000;1(1):31–9.

16. Lingwood D, Kaiser HJ, Levental I, Simons K. Lipid rafts as functional heterogeneity in cell membranes. Biochem Soc Trans. 21 sept 2009;37(5):955–60.

17. Lenne PF, Wawrezinieck L, Conchonaud F, Wurtz O, Boned A, Guo XJ, et al. Dynamic molecular confinement in the plasma membrane by microdomains and the cytoskeleton meshwork. EMBO J. 26 juill 2006;25(14):3245–56.

18. Lingwood D, Simons K. Lipid Rafts As a Membrane-Organizing Principle. Science. janv 2010;327(5961):46–50.

19. Pralle A, Keller P, Florin EL, Simons K, Hörber JKH. Sphingolipid–Cholesterol Rafts Diffuse as Small Entities in the Plasma Membrane of Mammalian Cells. J Cell Biol. 6 mars 2000;148(5):997–1008.

20. Sharma P, Varma R, Sarasij RC, Ira, Gousset K, Krishnamoorthy G, et al. Nanoscale Organization of Multiple GPI-Anchored Proteins in Living Cell Membranes. Cell. févr 2004;116(4):577–89.

21. Pike LJ, Casey L. Cholesterol Levels Modulate EGF Receptor-Mediated Signaling by Altering Receptor Function and Trafficking. Biochemistry. 1 août 2002;41(32):10315–22.

22. Low-Nam ST, Lidke KA, Cutler PJ, Roovers RC, van Bergen en Henegouwen PMP, Wilson BS, et al. ErbB1 dimerization is promoted by domain co-confinement and stabilized by ligand binding. Nat Struct Mol Biol. nov 2011;18(11):1244–9.

23. Bag N, Huang S, Wohland T. Plasma Membrane Organization of Epidermal Growth Factor Receptor in Resting and Ligand-Bound States. Biophys J. 3 nov 2015;109(9):1925–36.

24. Masson JB, Casanova D, Türkcan S, Voisinne G, Popoff MR, Vergassola M, et al. Inferring Maps of Forces inside Cell Membrane Microdomains. Phys Rev Lett. 29 janv 2009;102(4):048103.

25. Türkcan S, Masson JB, Casanova D, Mialon G, Gacoin T, Boilot JP, et al. Observing the Confinement Potential of Bacterial Pore-Forming Toxin Receptors Inside Rafts with Nonblinking Eu3+-Doped Oxide Nanoparticles. Biophys J. mai 2012;102(10):2299–308.

26. Türkcan S, Richly MU, Alexandrou A, Masson JB. Probing Membrane Protein Interactions with Their Lipid Raft Environment Using Single-Molecule Tracking and Bayesian Inference Analysis. Kanzaki M, éditeur. PLoS ONE. 3 janv 2013;8(1):e53073.

27. Türkcan S, Alexandrou A, Masson JB. A Bayesian Inference Scheme to Extract Diffusivity and Potential Fields from Confined Single-Molecule Trajectories. Biophys J. 16 mai 2012;102(10):2288–98.

28. El Beheiry M, Türkcan S, Richly MU, Triller A, Alexandrou A, Dahan M, et al. A Primer on the Bayesian Approach to High-Density Single-Molecule Trajectories Analysis. Biophys J. mars 2016;110(6):1209–15.

29. Floderer C, Masson JB, Boilley E, Georgeault S, Merida P, El Beheiry M, et al. Single molecule localisation microscopy reveals how HIV-1 Gag proteins sense membrane virus assembly sites in living host CD4 T cells. Sci Rep. déc 2018;8(1):16283.

30. Masson JB, Dionne P, Salvatico C, Renner M, Specht CG, Triller A, et al. Mapping the Energy and Diffusion Landscapes of Membrane Proteins at the Cell Surface Using High-Density Single-Molecule Imaging and Bayesian Inference: Application to the Multiscale Dynamics of Glycine Receptors in the Neuronal Membrane. Biophys J. janv 2014;106(1):74–83.

31. Tao RH, Maruyama IN. All EGF(ErbB) receptors have preformed homo- and heterodimeric structures in living cells. J Cell Sci. 1 oct 2008;121(19):3207–17.

32. Arkhipov A, Shan Y, Das R, Endres NF, Eastwood MP, Wemmer DE, et al. Architecture and Membrane Interactions of the EGF Receptor. Cell. janv 2013;152(3):557–69.

33. den Hartigh JC, van Bergen en Henegouwen PM, Verkleij AJ, Boonstra J. The EGF receptor is an actin-binding protein. J Cell Biol. 15 oct 1992;119(2):349–55.

34. Popoff. Epsilon toxin: a fascinating pore-forming toxin. FEBS J. 2011;278:4602.

35. Chakravorty A, Awad M, Cheung J, Hiscox T, Lyras D, Rood J. The Pore-Forming α-Toxin from Clostridium septicum Activates the MAPK Pathway in a Ras-c-Raf-Dependent and Independent Manner. Toxins. 10 févr 2015;7(2):516–34.

36. Newman, Schneider, Sutherland, Vodinelich, Greaves. The transferrin receptor. Trends Biochem Sci. 1982;7(11).

37. Srinivasan S, Regmi R, Lin X, Dreyer CA, Chen X, Quinn SD, et al. Ligand-induced transmembrane conformational coupling in monomeric EGFR. Nat Commun. 6 juill 2022;13(1):3709.

38. Iancu G, Serban D, Badiu CD, Tanasescu C, Tudosie MS, Tudor C, et al. Tyrosine kinase inhibitors in breast cancer (Review). Exp Ther Med. févr 2022;23(2):114.

39. Ivie SE, McClain MS. Identification of amino acids important for binding of Clostridium perfringens epsilon toxin to host cells and to HAVCR1. Biochemistry. 25 sept 2012;51(38):7588–95.

40. Santiago C, Ballesteros A, Tami C, Martínez-Muñoz L, Kaplan GG, Casasnovas JM. Structures of T Cell Immunoglobulin Mucin Receptors 1 and 2 Reveal Mechanisms for Regulation of Immune Responses by the TIM Receptor Family. Immunity. mars 2007;26(3):299–310.

41. Mukamoto M, Kimura R, Hang’ombe MB, Kohda T, Kozaki S. Analysis of tryptophan-rich region in Clostridium septicum alpha-toxin involved with binding to glycosylphosphatidylinositol-anchored proteins. Microbiol Immunol. 2013;57(3):163–9.

42. Hiroshima M, Abe M, Tomishige N, Hullin-Matsuda F, Makino A, Ueda M, et al. Membrane cholesterol interferes with tyrosine phosphorylation but facilitates the clustering and signal transduction of EGFR [Internet]. Biophysics; 2021 août [cité 28 oct 2022]. Disponible sur: http://biorxiv.org/lookup/doi/10.1101/2021.08.28.457965

43. Levitt ES, Clark MJ, Jenkins PM, Martens JR, Traynor JR. Differential Effect of Membrane Cholesterol Removal on μ- and δ-Opioid Receptors. J Biol Chem. août 2009;284(33):22108–22.

44. Sako Y, Kusumi A. Barriers for lateral diffusion of transferrin receptor in the plasma membrane as characterized by receptor dragging by laser tweezers: fence versus tether. J Cell Biol. 15 juin 1995;129(6):1559–74.

45. Türkcan S, Masson JB. Bayesian Decision Tree for the Classification of the Mode of Motion in Single-Molecule Trajectories. Hamacher K, éditeur. PLoS ONE. 20 déc 2013;8(12):e82799.

46. Pike LJ, Han X, Gross RW. EGF Receptors Are Localized to Lipid Rafts that Contain a Balance of Inner and Outer Leaflet Lipids: A Shotgun Lipidomics Study. 2005;27.

47. van der Maaten LJP, Hinton GE. Visualizing High-Dimensional Data Using t-SNE. J Mach Learn Res. 2008;9(nov):2579–605.

48. Danglot L, Chaineau M, Dahan M, Gendron MC, Boggetto N, Perez F, et al. Role of TI-VAMP and CD82 in EGFR cell-surface dynamics and signaling. J Cell Sci. 1 mars 2010;123(5):723–35.

49. Ibach J, Radon Y, Gelléri M, Sonntag MH, Brunsveld L, Bastiaens PIH, et al. Single Particle Tracking Reveals that EGFR Signaling Activity Is Amplified in Clathrin-Coated Pits. PLOS ONE. 17 nov 2015;10(11):e0143162.

50. Zhuo D, Guan F. Ganglioside GM1 promotes contact inhibition of growth by regulating the localization of epidermal growth factor receptor from glycosphingolipid-enriched microdomain to caveolae. Cell Prolif [Internet]. juill 2019 [cité 26 oct 2022];52(4). Disponible sur: https://onlinelibrary.wiley.com/doi/10.1111/cpr.12639

51. Charrin S, Jouannet S, Boucheix C, Rubinstein E. Tetraspanins at a glance. J Cell Sci. 1 sept 2014;127(17):3641–8.

52. Needham SR, Roberts SK, Arkhipov A, Mysore VP, Tynan CJ, Zanetti-Domingues LC, et al. EGFR oligomerization organizes kinase-active dimers into competent signalling platforms. Nat Commun. 31 oct 2016;7(1):13307.

53. Loi D, Mossa S, Cugliandolo LF. Effective temperature of active matter. Phys Rev E. 12 mai 2008;77(5):051111.

54. Vogtt K, Jeworrek C, Garamus VM, Winter R. Microdomains in Lipid Vesicles: Structure and Distribution Assessed by Small-Angle Neutron Scattering. J Phys Chem B. 29 avr 2010;114(16):5643–8.

55. Neisch AL, Fehon RG. Ezrin, Radixin and Moesin: key regulators of membrane-cortex interactions and signaling. Curr Opin Cell Biol. août 2011;23(4):377–82.

56. Jain AK. Data clustering: 50 years beyond K-means. Pattern Recognit Lett. 1 juin 2010;31(8):651–66.

